# Contactless metabolism estimation of small animals using high-frequency millimeter-wave radar

**DOI:** 10.1101/2024.04.11.588816

**Authors:** Hiroaki Ono, Kiyomi Ishikawa, Ayaka Wataki, Shoko Fujino, Genshiro A. Sunagawa

## Abstract

Animals flexibly adapt to internal and external environmental changes by utilizing energy produced from oxygen as fuel. By non-invasively monitoring an animal’s oxygen consumption, it becomes possible to understand an individual’s metabolic state. Calorimeters are known for directly measuring oxygen consumption but come with the issue of high initial costs. Despite the development of non-invasive techniques for measuring vital signs—including respiration rate (RR), heart rate (HR), and body temperature (Tb)—as indicators of metabolism, conventional methods encounter difficulties in estimating oxygen consumption rate (VO2). In this study, we developed a system that estimates the oxygen consumption of small animals using signals obtained from millimeter wave (mm-wave) radar technology processed through machine learning. By identifying frequency bands within the mm-wave signals contributing to VO2, our system is capable of predicting oxygen consumption several minutes in advance. Our system enables contactless, low-cost and multiplexed measurements of oxygen consumption, presenting a significant advancement in the field.

## INTRODUCTION

Energy balance is the ultimate boundary condition for living organisms. Animals expend finite energy resources to adapt to dynamic changes in their internal and external environments. For instance, animal survival is sustained by allocating finite energy to energy-consuming processes such as behavior, thermoregulation, and reproductive activities. Fundamental principles of energy production in living organisms are highly conserved across all life forms. Metabolism is the network of biochemical reactions employed by animals to transform molecules for the purpose of generating free energy (e.g., ATPs) and structural building blocks.

Adaptability is essential for metabolism to respond to changing internal and external environments. In vertebrates, the well-developed respiratory and cardiovascular systems enable a rapid adaptation to meet metabolic demands. For instance, the oxygen consumption rate (*VO_2_*) in mice increases linearly with work intensity. To match the elevated energetic demand, ventilation would be augmented at the transition from rest to running exercise, resulting in a 3-fold increase in RR^1–4^. Additionally, mice increase their HR during strenuous exercise, typically by ∼10 to 20%^5^. Conversely, in situations where oxygen demand is reduced, such as during hibernation, the respiratory and cardiovascular systems are suppressed. For a small mammal like the thirteen-lined ground squirrel (*Spermophilus tridecemlineatus*), physiological extremes reached during torpor include complete immobility, oxygen consumption maintained at 2 to 3% of the aroused condition, and HRs of 3 to 10 beats/min, compared with 200 to 400 beats/min when the animal is active^6^. Respiration is reduced from 100 to 200 to 4 to 6 breaths/min, and some species exhibit extended apneic periods^7^. In summary, metabolic demand reflects the overall state of an animal, and physiological parameters such as those from the respiratory and cardiovascular systems provide indirect insights into the overall metabolic demand by quantifying the metabolic rate.

Various technologies are currently employed for non-invasive monitoring of metabolic rate. The assessment of energy expenditure via direct/indirect calorimetry represents the most established methodology. Direct calorimetry involves the direct quantification of the body’s heat dissipation within a calorimeter chamber^8–10^. While offering high reproducibility and measurement errors ranging from 1–3%, the instrument is extremely expensive to build and run^8^. Indirect calorimetry, on the other hand, calculates energy expenditure based on the amounts of oxygen consumed and carbon dioxide produced. The most prevalent types of indirect calorimeters are ventilated, open-circuit systems, wherein the subjects are housed in airtight metabolic chambers with a continuous flow of fresh air^8,11^. The system collects and homogenizes the expired air, measures the flow rate, and analyzes oxygen and carbon dioxide concentrations in the incoming and outgoing air streams. These calorimeters are associated with substantial costs and exhibit slow response times. As discussed, the ability to measure respiratory rate (RR) and HR in a contactless manner can provide an indication of an animal’s metabolic state. Possible solutions for such contactless monitoring have been proposed through the utilization of video recording techniques. By leveraging video footage, algorithms can be employed to extract valuable physiological parameters, including RR and HR, without the need for direct physical contact or invasive procedures^12–15^. While video recording techniques have been proposed as potential solutions for contactless monitoring of metabolic indicators, they currently have significant limitations. The accuracy of HR detected from video footage remains questionable. Additionally, these video monitoring systems lose detection capability when mice are obscured by enrichment materials such as nesting pads within their housing environment. Recent advancements in mm-wave radar technology have led to its widespread use for vital sign monitoring, particularly in routine health monitoring applications^16–19^. The unique ability of mm-waves to penetrate clothing and body hair allows radar signals to capture movements associated with heat transfer and lung ventilation. This makes radar technology one of the most valuable methods for monitoring RR, HR, and muscle movements in a contactless manner^19–23^. While several technologies mentioned above can effectively extract vital sign signals, such as RR and HR, no studies have successfully estimated metabolic rates like *VO_2_*. This could be attributed to the necessity for nonlinear transformations to estimate *VO_2_* from vital signs or the need to integrate multimodal information such as ambient temperature (*Ta*) and body size, in addition to vital signs^24,25^.

Overall, contactless measurement of metabolic demands in animals is an ongoing challenge in metabolism research. In this study, we established an experimental framework that simultaneously recorded time-series mm-wave radar signals and *VO_2_* of small animals. By applying machine learning techniques to features extracted from the power spectrum (PS) of the mm-wave radar signals, we developed an algorithm capable of predicting instantaneous *VO_2_* from the radar signals and further forecasting future *VO_2_*. Furthermore, our approach successfully identified informative features from the mm-wave signals that correlated with metabolic expenditures. In summary, we present a novel machine learning-based algorithm for contactless estimation of metabolic expenditure in small animals using mm-wave radar technology.

## RESULTS

### Mm-wave radar system for recording the metabolism of free-moving small animals

We used a custom-made frequency-modulated continuous-wave (FMCW) radar system with a linear antenna array (Figure 1A). The FMCW radar emits a continuous wave signal whose frequency changes over time. This signal is reflected off a target and returns to the radar. There is a time delay and phase shift between the emitted signal and the signal that returns after reflection. Time delay and phase shift are directly related to the distance to the target and target velocity, respectively. Since the signal from the mm-wave radar should contain information on respiration, heartbeat, and muscle movements, we sought to use mm-wave radar signals to predict the metabolism of mice. The ground-truth metabolism was obtained by measuring mouse *VO_2_* by indirect calorimetry^11^. Mice were housed in temperature-controlled animal chambers, and the *VO_2_* was recorded by gas mass spectrometry every 6 minutes. At the same time, we set up mm-wave radars that were installed 40 cm away from the chambers (Figure 1B). The mm-wave radar generated complex-valued time-and-distance dependent signals *S_c_ (t, r)* at a 100 Hz sampling rate. Because the variance of mm-wave signals reflected from objects moving like mice should be greater than that of signals reflected from stationary objects such as walls, we manually select *r* with a relatively higher variance that should correspond to the position of the target subject (Figure S1). In this setting, we have 36,000 data points from mm-wave radar signals per single distance channel to predict a single data point of *VO_2_* (Figure 1C). We used the inbred mouse strain C57BL/6J to observe daily torpor during measurement periods, which stably enters torpor under certain conditions. By selecting male mice aged between 7 and 9 weeks, our goal was to minimize phenotypic variance. Mice were subjected to recording for more than three days in a constant *Ta*. At the beginning of the second day, food was removed for 24 hours. A typical torpor episode began on the latter half of the same day (Figure 1D). We successfully recorded the animals’ *VO_2_* and mm-wave signals *S_c_ (t, r)* simultaneously without contact (Figures 1E, S2–3). *VO_2_* exhibits a higher median value at lower *Ta*s, underscoring the higher oxygen demand to maintain *Tb* in environments cooler than the thermoneutral zone. Specifically, at a *Ta* of 20 °C, *VO_2_* stands at 2.877 ml/g/h with a standard deviation of 0.606 ml/g/h, based on sample size of n = 4; at 28°C, it decreases to 1.792 ± 0.555 ml/g/h (n = 4); and at 12°C, it rises to 3.459 ± 0.661 ml/g/h (n = 4) (Figure S2).

**Figure 1.**
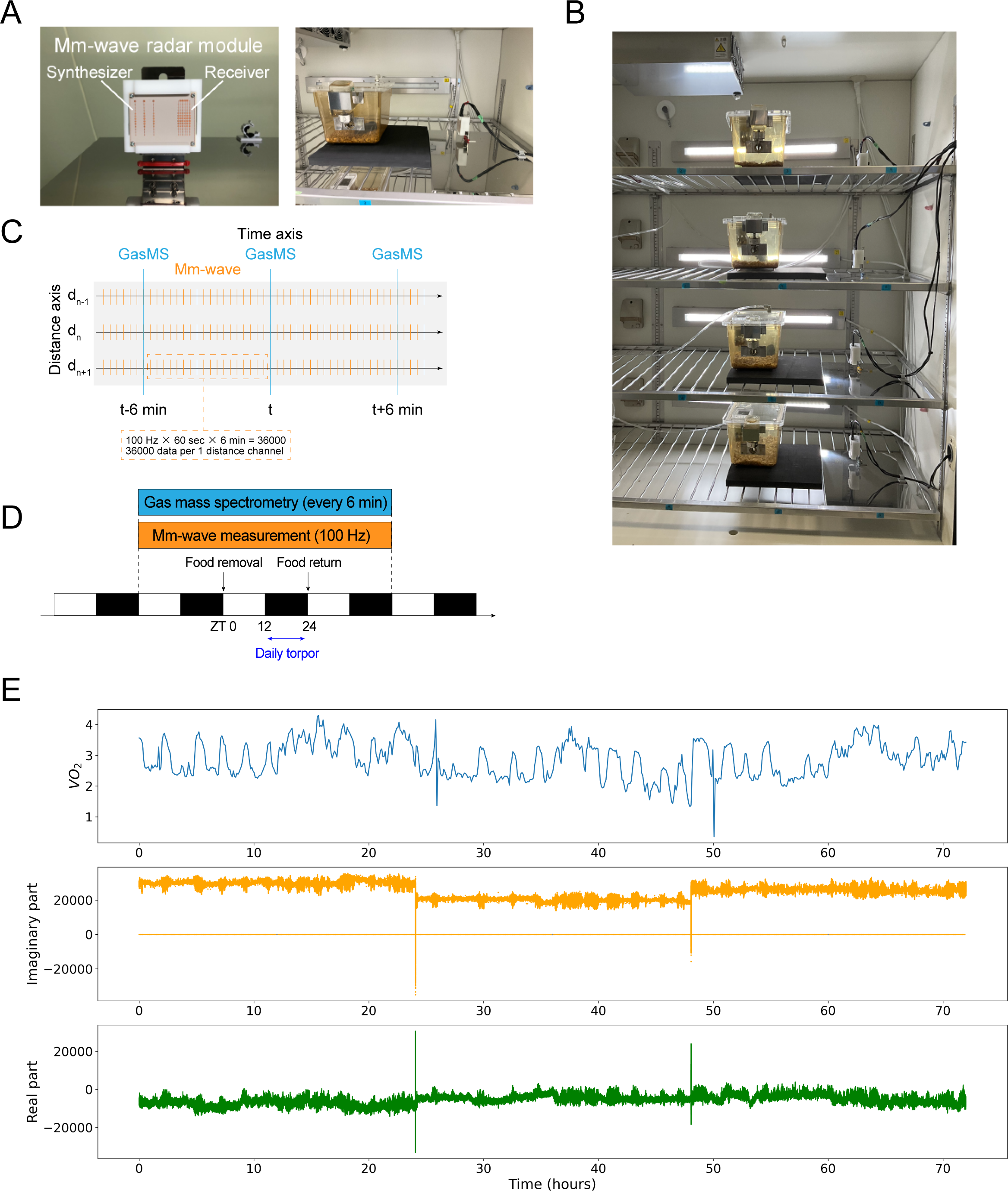
Experimental setup for metabolic measurement using mm-wave radar technology. (A) Overview of a mm-wave radar module. With dimensions of 6.5 cm by 3.2 cm by 6.1 cm, the module is compact enough to be installed in a standard mouse experimental setting. (B) A system designed for measuring mm-wave radar signals combined with evaluating the metabolism of freely moving mice^11^. The temperature-controlled animal chamber (left panel) and the inside of the chamber (middle panel), in which mm-wave radar was installed 40 cm away from a metabolic chamber, are shown. The *VO_2_* is recorded by gas mass spectrometry. € Scheme of animal experiments. (D) Scheme of measurement conditions for mm-wave radar signals and *VO_2_*. (E) Example plot of simultaneously recorded *VO_2_* and mm-wave radar signals, which include both imaginary and real components.

### Metabolism estimation using machine learning-based mm-wave radar technology

Because the mm-wave radar sampled signals at the rate of 100 Hz, the signals would contain information on RR, HR, and skeletal muscle movements, which have slower changes than the sampling rate. We used this information to predict the *VO_2_* from the mm-wave radar signal every six minutes. By applying the short-time Fourier transform (STFT) to six-minute segments of these radar signals, we generated spectrograms to extract features from the frequency domain. The mm-wave spectrogram is a 256 × 283 pixel two-dimensional histogram with time and frequency axes and values colored by their power. After reshaping the STFT data into a resolution of 224 × 224 pixels by a bilinear interpolation algorithm, we standardized the spectrograms by subtracting the global mean pixel value of the entire dataset of a mouse from each individual pixel value and then dividing by the global standard deviation (Figures 2A and 2B). We treated the spectrograms as image data, allowing us to apply well-developed pre-trained models for image recognition with some modifications. We constructed the machine learning model using CNNs based on the EfficientNetV2-B0 model, developed originally for classifying a large set of images (ImageNet)^26^. Our model was based on an ensemble learning approach consisting of two transfer learning models (Figure 2C). Both models use three spectrogram images from time *t* to *t-2* as pseudo-3-channel RGB. The *Ta* was also added to the input.

**Figure 2.**
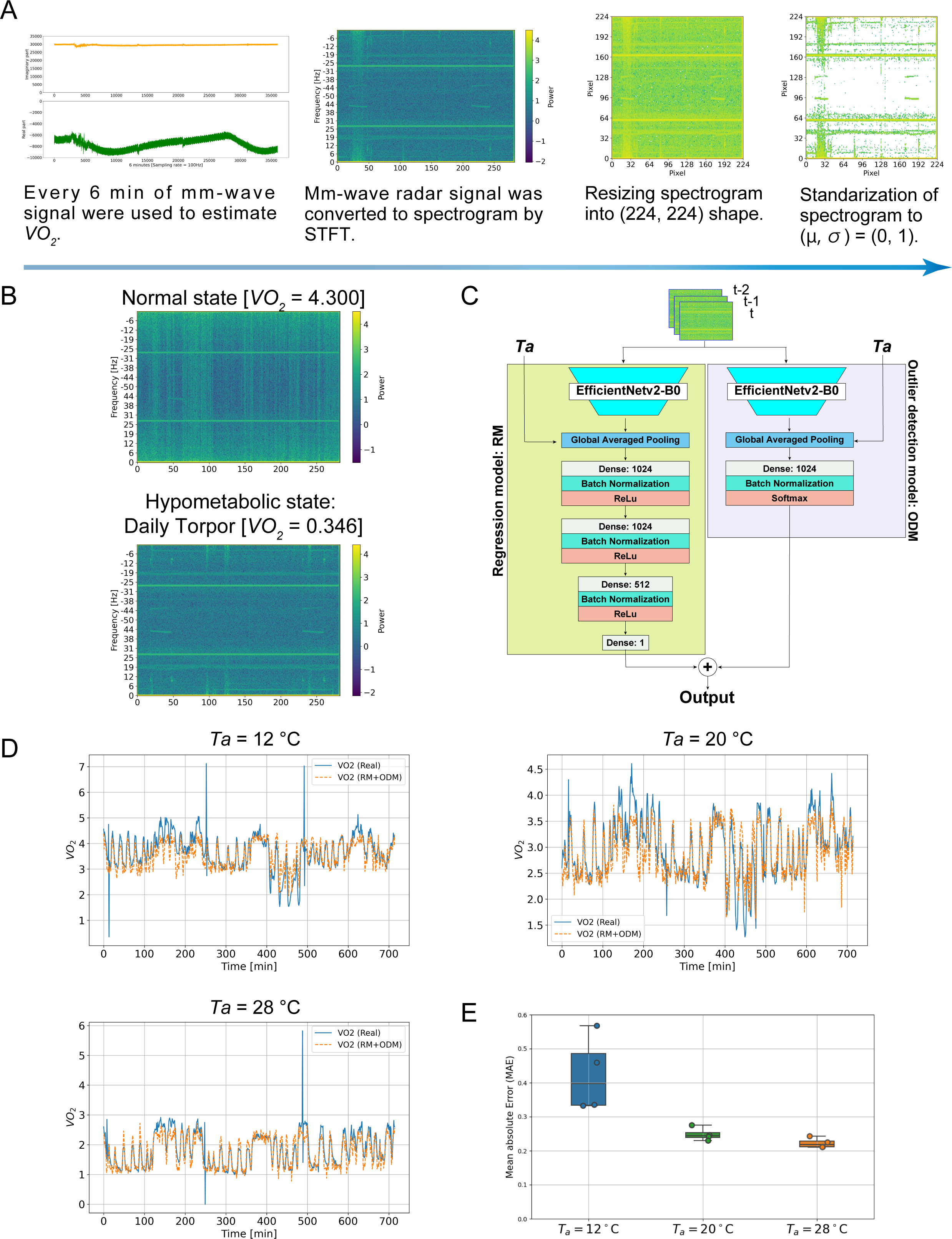
Metabolism estimation using machine learning-based mm-wave radar technology. (A) The mm-wave radar signal preprocessing pipeline. For each 6-minute interval of *VO_2_* measurement, 36,000 points of the imaginary and real components were utilized. First, the imaginary and real parts were combined into a complex-valued time series data, which was then converted into a power spectrogram using a Short-Time Fourier Transform (STFT) with a window size of 254. In the following processes, the components of the negative frequencies were included in the computation. Next, the spectrogram was resized to a shape of (224, 224), after which it was standardized to have a mean of 0 and a variance of 1. (B) Examples of heatmaps obtained from a normal state (*VO_2_* = 4.300 ml/g/h) and a hypometabolic state (daily torpor, *VO_2_* = 0.346 ml/g/h). (C) A schematic diagram of the ensembled transfer learning with temporal convolution. To regress *VO_2_* at a given time t, the spectrograms obtained from the three most recent time frames (t-2 to t) were used as pseudo-3-channel RGB images. The RM starts with the convolutional layers of EfficientNetV2-B0, followed by three dense layers (1024, 1024, 512 units) with the identity function as the final activation. The ODM also starts with the EfficientNetV2-B0’s convolutional layers but followed by a single dense layer. For *VO_2_* categorization into ‘Normal’, ‘Lower’, or ‘Higher’, the final activation used was the softmax function. The outputs from both models underwent a linear transformation: if ODM categorized the output as ‘Lower’, RM’s prediction was scaled down by 25%; if ‘Higher’, it was increased by 10%. (D) Examples of time-series data displaying predicted *VO_2_* values and actual *VO_2_* values. The blue line represents the actual *VO_2_* values, while the orange dashed line indicates the predicted *VO_2_* values. (E) The performance of MMM at different *Ta*s.

The Regression Model (RM) is designed to regress oxygen consumption rate from three spectrograms. These spectrograms are input into the convolutional layers of the EfficientNetV2-B0 as three-channel image data. Then, they are concatenated in the global average pooling layer with *Ta* as numerical data. Three fully connected layers with 1024, 1024, 512, and 1 output layer were added. The activation function was rectified linear function (ReLU), and batch normalization was added between the fully connected layers to avoid overfitting.

The Outlier Detection Model (ODM) is designed to predict outlying *VO_2_* values, defined as values deviating by more than ±1 ml/g/h from each mouse’s median *VO_2_*. Like the RM, it consists of convolutional layers from the EfficientNetV2-B0 and fully connected layers, but its final activation function is the softmax function. The outputs are then classified into three classes: typical values (Normal *VO_2_*), values lower than the median (Lower *VO_2_*), and values higher than the median (Higher *VO_2_*).

The final output was generated by integrating the RM and ODM outputs. Specifically, if the ODM classifies the input as ‘Normal *VO_2_’*, the output of RM was retained without modification (multiplied by 1). For inputs classified as ‘Lower *VO_2_’*, the RM output was multiplied by 0.75, reflecting a 25% reduction. Convers*VO_2_*ely, for ‘Higher *VO_2_*’ classifications, the RM output was increased by a factor of 1.1, indicating a 10% increase. This approach applied a linear transformation to the RM output, effectively calibrating the final output according to the *VO_2_* status identified by the ODM.

We measured *VO_2_* and mm-wave radar signals across various *Ta*s of 12 °C, 20 °C, and 28 °C, utilizing the protocol mentioned above (Figure 1D). These measurements were carried out on groups of four mice for each *Ta* setting. A recording from one mouse was set aside as test data, and the remaining data from 11 mice were randomly split into training data (eight mice) and validation data (three mice). We optimized each model, RM, and ODM, by fine-tuning the parameters across all layers. We evaluated the model architecture for each *Ta* using mean absolute errors (MAEs) as a metric. Following the Leave-One-Out Cross-Validation method, we found that our model achieved a MAE of 0.244 ± 0.0167 ml/g/h at *Ta* = 20 °C, 0.218 ± 0.0130 ml/g/h at *Ta* = 28 °C, and 0.397 ± 0.0976 ml/g/h at *Ta* = 12 °C.

While MAE at 12 °C is relatively high, we found that our models could predict features of the diurnal rhythm and daily torpor in the *VO_2_* from the mm-wave radar signals across all *Ta*s (Figures 2D, 2E, and S4). In this study, we named the system ‘Machine-learning-based Metabolism Estimation using Millimeter-wave Radar (MMM).’

### Visualization of informative features from intermediate layers

To make deep learning decoding interpretable, we tried to quantify the contribution of the spectrogram to the regression in the final output from the convolutional layers of RM. To achieve this, we used a visualization technique known as Gradient-Class Activation Mapping (Grad-CAM)^27^. In brief, Grad-CAM first computes the gradient of the predicted values with respect to feature map activations of a last convolutional layer. By visualizing these gradients as heat maps, we can gain insights into which feature map activations contribute most significantly to the predicted values. We obtained visual explanations of the feature map activation (7 × 7 in shape) in the final layer of the RM using Grad-CAM. Subsequently, we flattened the feature map activations to 49 x 1, grouped them by Ta, and performed principal component analysis (PCA) (Figure 3A). As a result, while no axis reflecting the *VO_2_* feature could be found at *Ta* = 12 °C, a *VO_2_* gradient was observed along the principal component 1 (PC1) axis at *Ta* = 28 °C and along the PC2 axis at *Ta* = 20 °C (Figures S5 and 3B). In *Ta* = 28 °C with high predictive performance, MMM accurately extracts *VO_2_* information from the spectrogram. Therefore, we chose *Ta* = 28 °C for the following analysis to understand the explanatory ability of the MMM. We calculated the loading of the PC1 component at *Ta* = 28 °C for each pixel, resized it to the same shape as the input spectrogram (224 × 224) (Figure 3C). As a result, the loading was high for both positive and negative high-frequency components on the frequency axis and around 3 minutes on the time axis. It can be considered that *VO_2_* is represented in the certain dimensions that have specific features.

**Figure 3.**
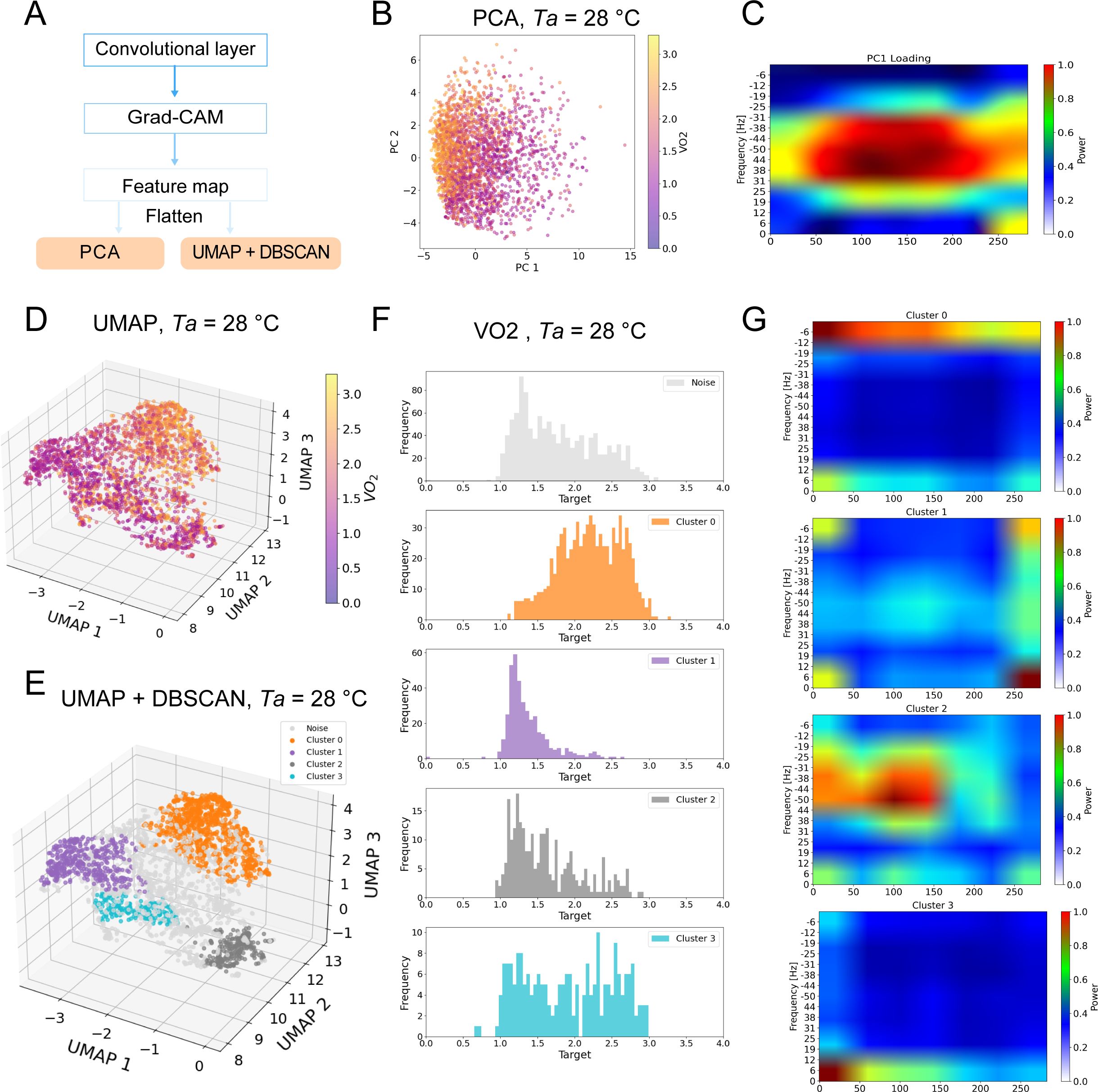
Visualization of informative features in MMM. (A) The workflow for visualizing explanations in MMM. (B) The PCA result at *Ta* = 28 °C. The color gradient represents the *VO_2_* value at each point. (C) Visualization of the first principal component (PC1) loadings in spectrogram format (283 × 256 pixels). (D) UMAP dimensionality reduction at *Ta* = 28 °C. Similar to PCA, the color gradient represents the *VO_2_* value at each point. (E) UMAP dimensionality reduction followed by DBSCAN clustering. Excluding noise points, the data was divided into four distinct clusters named cluster 0 to cluster 3. (F) Distribution of *VO_2_* values within each cluster. (G) Visualization of explanations within each cluster in a spectrogram format (283 × 256 pixels).

To understand the dimensions that is meaningful for the regression of *VO_2_*, we applied dimension reduction using a different method, uniform manifold approximation and projection (UMAP). The 49 feature pixels obtained by Grad-CAM were visualized in a 3-dimensional space using UMAP (Figure 3D). We applied the clustering method known as density-based spatial clustering of applications with noise (DBSCAN) to the feature map activation (Figure 3E). As a result, we obtained four clusters (clusters 0 to 3) as shown in Figure 3D. The distribution of *VO_2_* in each cluster is shown in Figure 3E. Cluster 0 is most frequent and has high *VO_2_*. Cluster 1 shows the lowest *VO_2_*, suggesting that the feature map activation during a hypometabolic state such as daily torpor is included. Clusters 3 and 4 are clusters between the normal metabolic and hypometabolic states. Since cluster 4 has peaks in both the high and low *VO_2_* parts, it is considered that cluster 4 is closer to cluster 1 than cluster 3 in the state space of the metabolic state. Finally, to visualize the explanation of the MMM in each cluster, we averaged the feature map activation for each cluster, resized it to the shape of the spectrogram, and plotted it. As a result, it was found that there are characteristic patterns in both positive and negative frequency bands for each cluster for the regression of *VO_2_*. The explanation lies in the low-frequency component over the entire time in clusters containing relatively high VO2 (clusters 0 and 3). The explanation for relatively low *VO_2_* (clusters 1 and 2) lies in the high-frequency component at intermediate times (3 min). In particular, it was found that cluster 1 has the most explanatory components in the negative low-frequency area in the latter half of the time in addition to the high-frequency component.

Thus far, it has been found that the information on *VO_2_* is embedded in both positive and negative frequencies in the spectrogram. The MMM can regress *VO_2_* by extracting this information.

### Performance evaluation of future metabolic state

Given that MMM processes consecutive 18-minute segment data as input, it potentially captures local trends and distributions in the time-series data of oxygen consumption rate. Furthermore, the mm-wave signal contains real-time physiological information, such as HR, RR, and skeletal muscle movement. It is plausible that *VO_2_* attributed to these activities may be discernible with a certain time delay. To verify whether MMM could predict the future *VO_2_*, we trained the MMM parameters by shifting the *VO_2_* values by 6*t* minutes (*t* = 1, 2, 3, 4, 5, and 10) into the future (Figure 4A). To estimate the worst or the upper bound of the MAE in this model, we trained the model with a training and validation dataset where the *VO_2_* values were randomly shuffled within each mouse (Figure 4B). We found that the upper bound of MAE of 0.525 ± 0.00589 ml/g/h at 20 °C, 0.532 ± 0.0482 ml/g/h at 28 °C, and 0.555 ± 0.0269 ml/g/h at 12°C. For all *Ta* conditions, the MAE gradually increases in proportion to the length of the forecast period for the *VO_2_*. Particularly at 12 °C, the MAE rises quickly, reaching the random level when predicting more than 12 minutes into the future (Figure 4C). In contrast, for 20 °C and 28 °C, MAEs increase relatively gradually (Figures 4D and 4E). In both cases, when predicting 60 minutes into the future, the MAE reaches the same level as the random case. Particularly at 28°C, we found that even when predicting 18 minutes into the future, the MAE remained low enough that daily torpor could be detected (Figure 4E).

**Figure 4.**
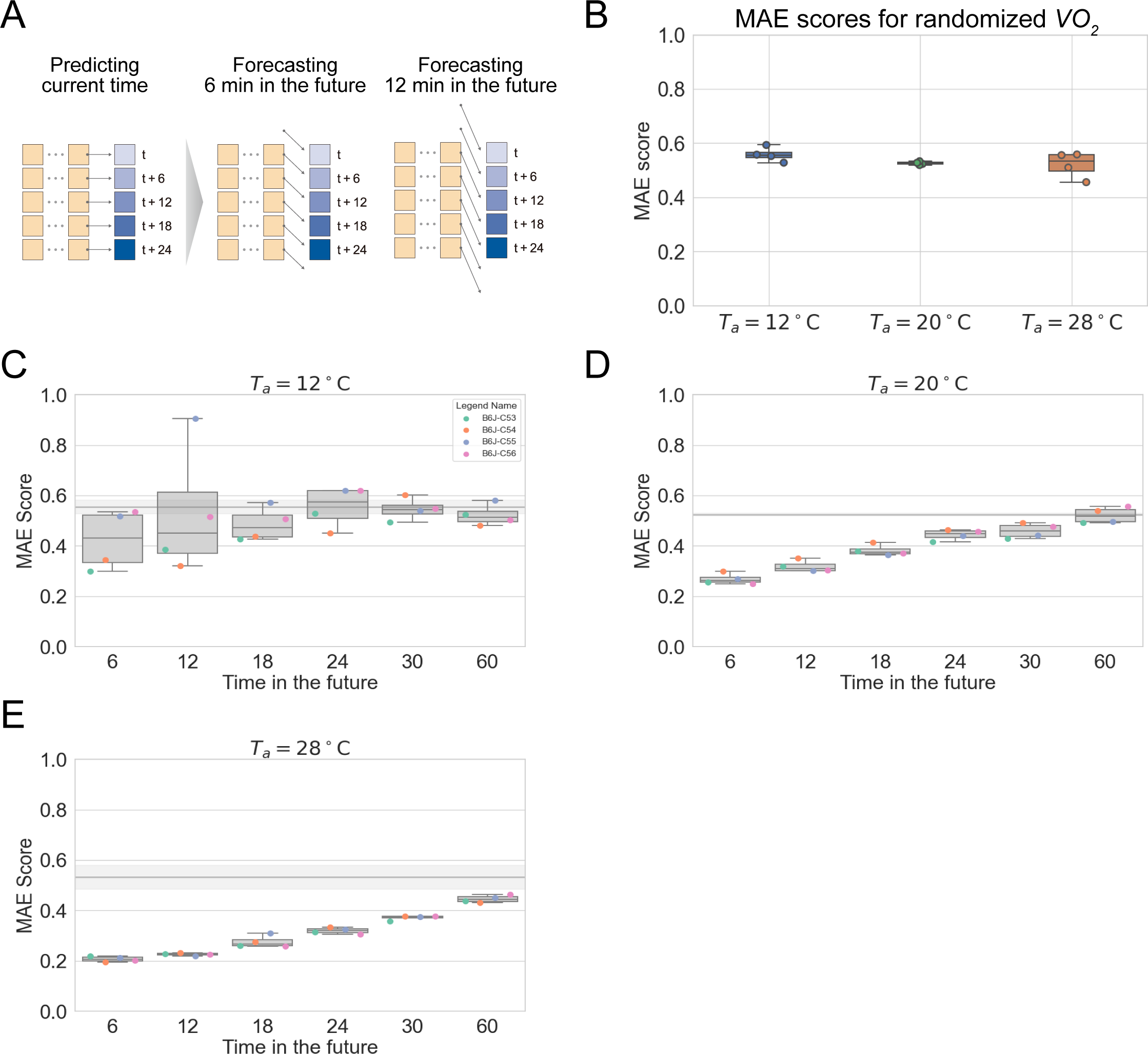
Performance evaluation of future metabolic state. (A) The experimental scheme for forecasting future *VO_2_* values. During the training phase, MMM was fed by time-shifted mm-wave radar signals and corresponding *VO_2_* values. This transformed the task into forecasting *VO_2_* at a future time point determined by the shift duration, with each shift corresponding to a 6-minute interval. For example, shifting the *VO_2_* data by one time step resulted in an MMM model that predicted *VO_2_* 6 minutes ahead, while a two-step shift yielded a model forecasting *VO_2_* 12 minutes into the future. (B) MAEs for each *Ta* when the *VO_2_* values are shuffled randomly. (C–E) Box-and-whisker plots depicting the change in MAE scores when forecasting *VO_2_* up to 60 minutes into the future for each *Ta*. The gray line and shaded region represent the median and standard deviation of the MAE obtained when predicting randomly shuffled *VO_2_* values in Figure 4B (*Ta* = 12 °C, 0.555 ± 0.0269 ml/g/h; *Ta* = 20 °C, 0.525 ± 0.006 ml/g/h; *Ta* = 28 °C, 0.533 ± 0.048 ml/g/h).

## DISCUSSION

### Mm-wave radar-based monitoring with machine learning provides the prediction of oxygen consumption rate

Metabolism represents a dynamic state, readily influenced by circadian rhythms, sleep-wake cycles, seasons, rearing environments, experimental manipulations, and various other factors^28–32^. Therefore, to study the dynamics and adaptability of metabolism, minimizing the external effect on the metabolism in a contactless manner is mandatory. Numerous methods have been developed to measure metabolic parameters without any physical contact. For example, video-based vital monitoring^12–14^ is one of the major methods for estimating metabolism, as vital signs directly reflect metabolic adaptation. Another approach is to use thermography^33,34^, given that a significant amount of oxygen is used in the production of heat to maintain *Tb* in homeotherms, especially in small mammals^35^. Capacitive sensing-based method is also developed^36^. Recently, various radar-based technologies to track physiological status have been developed ^37–39^. Among them, the mm-wave radars are one of the most promising approaches that enable contactless monitoring of vital signs. Mm-wave radar allows measuring distances and displacements with sub-mm precision, sufficient to capture the small displacements caused by respiration and heart activity. Many mm-wave radar-based vital sensing methods have been developed, especially for humans. Previous studies estimated the HR and RR from the analysis of displacement signals by conventional signal processing techniques like Fast Fourier Transform (FFT)^40^. In addition, previous work has also used empirical mode decomposition (EMD)^41^, Random Body Motion Cancellation algorithms^42^ or feature-based approach^22^ to estimate HR more precisely. Although the methods mentioned thus far can accurately detect HR, none can estimate *VO_2_* in vivo.

In response to this limitation, we developed a machine learning–based model that estimates *VO_2_* from mm-wave radar signals, bypassing the need for RR or HR measurements. We demonstrated that median MAEs are less than 0.4 ml/g/h at *Ta* = 12, 20, and 28 °C (Figure 2). A unique aspect of our model lies in its utilization of time-series spectrogram data as input. The original mm-wave radar signal, inherently multi-dimensional time-series data, can transformed into spectrograms, effectively treating it as image data for analysis. Feeding CNN models with spectrograms has been applied in diverse applications, including hard X-ray flare analysis^43^, in vivo Ca^2+^ imaging^44^, and ECG analysis^45^. This approach of interpreting time-series data as images enables the application of the EfficientNetV2-B0 model^26^, which has been successful in image recognition competitions.

The MMM has higher MAE at *Ta* = 12 °C than at *Ta* = 20 °C or *Ta* = 28 °C. Since the thermoneutral zone for mice is ∼29–33 °C^46^, mice should feel cold at *Ta* = 12 °C resulting in heat production using oxygen^11^. In Figure 3, we demonstrated that the multiple clusters were separated for hypometabolism (clusters 1, 2, and 3), but cluster 0 was the only cluster for normal to hypermetabolism. This may cause relatively low predictive performance of the MMM in cases of high oxygen consumption rate, as in at *Ta* = 12 °C.

### Potential application of forecasting metabolic state

We can assume that *VO_2_* at a given time should be determined by integrating the recent energy consumption from every tissue. Moreover, if the animal shows a certain movement that mm-wave radars can detect before the metabolism changes, the future *VO_2_* estimation can be further improved. Therefore, in principle, we should be able to forecast *VO_2_* to some extent from the current mm-wave data, which includes metabolism data and the metabolism determinant movements. In Figure 4, we demonstrated the near future forecast is possible from current mm-wave radar signals. The condition where *VO_2_* was randomly shuffled can be considered a condition where there is no information about *VO_2_* in the spectrogram. Compared to this randomized condition, the mm-wave radar signals under *Ta* = 20 °C and *Ta* = 28 °C are considered to retain predictive metabolic information up to 60 minutes into the future. While the contributing factors remain unidentified, the current study strongly suggests the existence of a component characterized by slow dynamics, spanning several minutes, among the factors influencing *VO_2_*.

One area of research that would benefit from the MMM’s ability to predict near-future metabolic states is the exploration of factors that induce hibernation. To induce hibernation under laboratory conditions, a hibernator, such as Syrian golden hamster (*Mesocricetus auratus*), have to be housed under a short-day photoperiod and cold *Ta*s for several months^47^. At present, predicting when the animal enter hibernation is unfeasible. However, if MMM could predict the entry timing into hibernation a few minutes in advance, it would enable us to sample tissues that should contain the hibernation induction factors.

### Limitations of this study

Currently, MMM has not succeeded in prediction of *VO_2_* in real-time. In this study, the spectrogram is standardized based on the distribution information of three consecutive days of mm-wave radar signals. Therefore, to enable real-time prediction, the distribution of mm-wave radar signal prior to the recording is required. For example, obtaining the signal for three days beforehand would be necessary for real-time prediction scenarios.

## ACKNOWLEDGMENTS

We are grateful to members of the Sunagawa lab for their valuable discussions, insightful advice, and daily support with our research. Additionally, our gratitude extends to LARGE, RIKEN BDR for providing exceptional mouse housing facilities. This work was supported by Grant-in-Aid for Early-Career Scientists from JSPS (23K14130) (H.O.), the RIKEN BDR Quiescence Manipulation and Investigation of Natural-hypometabolism (QMIN) Project (G.A.S.), the Grant-in-Aid for Challenging Research (Pioneering) from JSPS (21K18276) (G.A.S.), the Grant-in-Aid for Transformative Research Areas (A) from JSPS (23H04941) (G.A.S.), and the Suntory Rising Stars Encouragement Program in Life Sciences (SunRiSE) (G.A.S.).

## AUTHOR CONTRIBUTIONS

H.O. and G.A.S. designed the study. H.O. did computational framework and analyzed data. K.I., A.W. and S.F. performed the animal experiment. H.O. and G.A.S. wrote the manuscript.

## INCLUSION AND DIVERSITY

We support inclusive, diverse, and equitable conduct of research.

## STAR★METHODS

### KEY RESOURCE TABLE

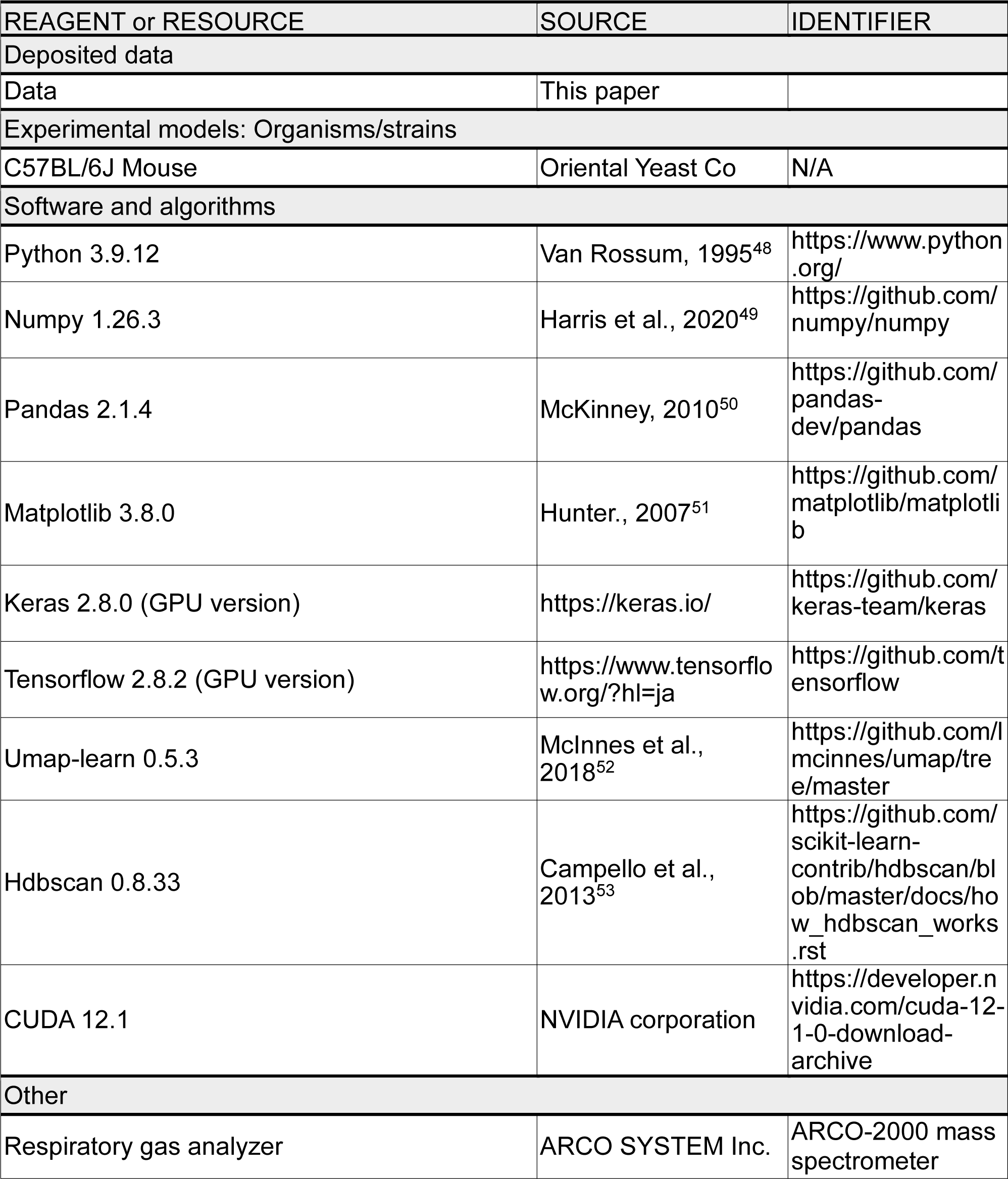

## RESOURCE AVAILABILITY

### Lead contact

Further information and requests for resources should be directed to and will be fulfilled by the lead contact, Genshiro A. Sunagawa.

### Materials availability

This study did not generate new unique reagents.

### Data and code availability

Any additional information including code reported in this paper is available for non-commercial use from the lead contact upon reasonable request.

## EXPERIMENTAL MODEL AND SUBJECT DETAILS

### Animals

C57BL/6J mice were purchased from Oriental Yeast Co. Until the mice were used in torpor experiments, they were given food and water ad libitum and maintained at a *Ta* of 21 °C, relative humidity of 50%, and a 12 h light/12 h dark cycle (8 a.m. lights on, 8 p.m. lights off). In the simultaneous recording of mm-wave radar signals and *VO_2_* (Figure 1), twelve 12-week-old C57BL/6J male mice were used. All animal experiments were approved by the Institutional Animal Care and Use Committee of RIKEN Kobe Branch. All animal experiments were performed according to RIKEN Regulations for Animal Experiments.

## METHOD DETAILS

### Oxygen consumption recordings

Each animal was housed in a temperature-controlled chamber (Cat# LP-400P-AR, Nippon Medical & Chemical Instruments Co., Ltd.). Oxygen consumption and carbon dioxide output rates (*VO_2_* and *VCO_2_*, respectively) of the animal were continuously recorded with a respiratory gas analyzer (ARCO-2000 mass spectrometer, ARCO SYSTEM Inc.). The respiratory coefficient was calculated as the ratio of VCO2 to *VO_2_*.

### Fasting-induced torpor

FIT induction experiment (Figure 1) was designed to record the animal’s metabolism for 3 days. The animals were introduced to the recording chamber the day before recording started (day 0). Food and water were freely accessible. *Ta* was set as indicated on day 0 and kept constant throughout the experiment. On day 2, ZT-0 (8 a.m.), the food was removed to induce torpor. After 24 h, on day 3, ZT-0 (8 a.m.), the food was returned to each animal chamber.

### Mm-wave radar recording

The mm-wave radar module (MaRI Co., Ltd.) was installed approximately 40 cm away from the metabolic chamber. The mice were housed in a temperature-controlled environment (*Ta* = 12, 20, and 28 °C). For each *Ta* condition, 4 mice were recorded. The mm-wave radar signal was recorded as complex numbers at a sampling frequency of 100 Hz and saved in Numpy format on the attached computer.

### Data allocation

Leave-One-Out Cross-Validation (LOOCV) was employed to assess the model’s performance and generalization ability. The dataset consisted of measurements from 12 mice. For each iteration of the LOOCV, one mouse was held out as the test set. The remaining 11 mice were then randomly permuted. From this permuted list, eight mice were randomly selected, and their corresponding data was concatenated to form the training set. The data from the remaining three mice was used as the validation set. The model was trained on the training set and evaluated on both the validation set and the test set, which was the held-out mouse. This process was repeated 12 times, with each mouse being used exactly once as the test set.

### Data and preprocessing

In the data preprocessing stage, several steps were performed to transform the input data into a suitable format for the deep learning model. First, the complex-valued input signals were converted into spectrograms using the Short-Time Fourier Transform (STFT), with a window size of 256 and a hop length of 128 and returned the absolute value of the complex STFT coefficients. The spectrograms were then resized to a fixed size of 224 × 224 pixels. Each frame of the spectrogram was resized individually using bilinear interpolation and then stacked the resized frames to create the final spectrogram. Additionally, a logarithmic operation was applied to the resized spectrograms. We standardized spectrograms for each mouse separately by subtracting the mean and dividing by the standard deviation of the data points corresponding to that mouse. Finally, a temporal convolution operation was applied to the standardized spectrograms. The data were divided into blocks of three consecutive frames and concatenated them along the channel dimension, effectively creating a 3-channel input for the deep learning model. This step was performed for each group of data points corresponding to a single mouse.

### Preparation of training dataset for ODM

For the mice allocated as the training dataset and the validation dataset, the *VO_2_* median for each mouse was used to define *VO_2_* levels: values greater than 1 above the median were classified as higher *VO_2_*, values more than 1 below as lower *VO_2_*, and values within a ±1 range of the median as normal *VO_2_*. Then, labels were assigned to each *VO_2_* level using one hot encoding.

### Deep learning

Deep learning was performed using Python 3.9.12, Anaconda Packages, Keras 2.8.0, Tensorflow 2.8.2 (https://www.tensorflow.org/?hl=ja). The RM architecture was based on the EfficientNetV2-B0 convolutional neural network (CNN), pre-trained on the ImageNet dataset^26^. Two input tensors were defined: one input with a shape of (224, 224, 3) for the time-series spectrogram, and another input with a shape of (1,) for the *Ta*. The EfficientNetV2-B0 model was loaded without the top layer, using the time-series spectrogram (224, 224, 3) as input. All layer of EfficientNetV2-B0 CNN was set to trainable, allowing the weights to be updated during training. A GlobalAveragePooling2D layer was applied to the output of the EfficientNetV2-B0 model to reduce the spatial dimensions. The output of the GlobalAveragePooling2D layer and the *Ta* input were concatenated. This concatenated tensor was then passed through a series of Dense layers with ReLU activation functions and BatchNormalization layers: a Dense layer with 1024 units and ReLU activation, followed by a BatchNormalization layer, another Dense layer with 1024 units and ReLU activation, another BatchNormalization layer, a Dense layer with 512 units and ReLU activation, another BatchNormalization layer, and finally, a Dense layer with 1 unit without an activation function. The model was compiled with the Adam optimizer, with a learning rate set to 0.00001. The mean squared error (MSE) was used as the loss function, and the mean absolute error (MAE) was included as an evaluation metric. The training was carried out for 300 epochs with a batch size of 32.

The ODM architecture was also based on the EfficientNetV2-B0 CNN loaded without the top layer. Two input tensors were defined: one for the image input with a shape of (224, 224, 3), and another for the *Ta* input with a shape of (1,). A GlobalAveragePooling2D layer reduced the spatial dimensions of the EfficientNetV2-B0 output, which was then concatenated with the *Ta* input. This tensor passed through Dense layers with ReLU activation and BatchNormalization. The final layer was a Dense layer with 3 units and softmax activation for 3-class classification. The model used the Adam optimizer with a learning rate of 0.0001, categorical cross-entropy loss, and accuracy metric. Before training, the labels were assigned weights. The weights were set to 1 for normal *VO_2_*, 10 for lower *VO_2_*, and 1.5 for higher *VO_2_*. The training was carried out for 25 epochs with a batch size of 32.

### Visual explanation of MMM as a heatmap

Gradient-weighted Class Activation Mapping (Grad-CAM) was employed to visualize the important regions in the input spectrograms that contributed to the model’s predictions^27^. Grad-CAM uses the gradients of the target prediction flowing into the final convolutional layer to generate a heatmap highlighting the important features learned by the model. A tensorflow gradientTape function was used to capture the gradients of the model’s output with respect to the output of the last convolutional layer. The gradients were then pooled across the spatial dimensions and used to compute the heatmap by taking the product with the output of the last convolutional layer. The size of heatmap was (7, 7) shape. For each time step in the test dataset, the heatmap for the corresponding input data (spectrogram and *Ta*) were computed.

### Dimension reduction using PCA or UMAP

The heatmap obtained from Grad-CAM, with a shape of (7, 7), was flattened into a shape of (49,). Each element of this flattened array was then used as a feature for performing Principal Component Analysis (PCA). The data was visualized in two dimensions along the PC1 and PC2 axes. UMAP constructs a high-dimensional graph representation of the data and then optimizes a low-dimensional graph to be as structurally similar as possible. We performed UMAP using the Umap-learn 0.5.3 library with the following parameters: n_components = 3, metric = ‘manhattan’, n_neighbors = 15, min_dist = 0.01. The data was visualized in a three-dimensional space along the UMAP1, UMAP2, and UMAP3 axes. We employed HDBSCAN to identify clusters within our dataset following dimensionality reduction via UMAP. We initialized the HDBSCAN clustering algorithm using the hdbscan package 0.8.33 with the following parameters: min_cluster_size = 50, min_samples = 80.

---

**Figure S1.**
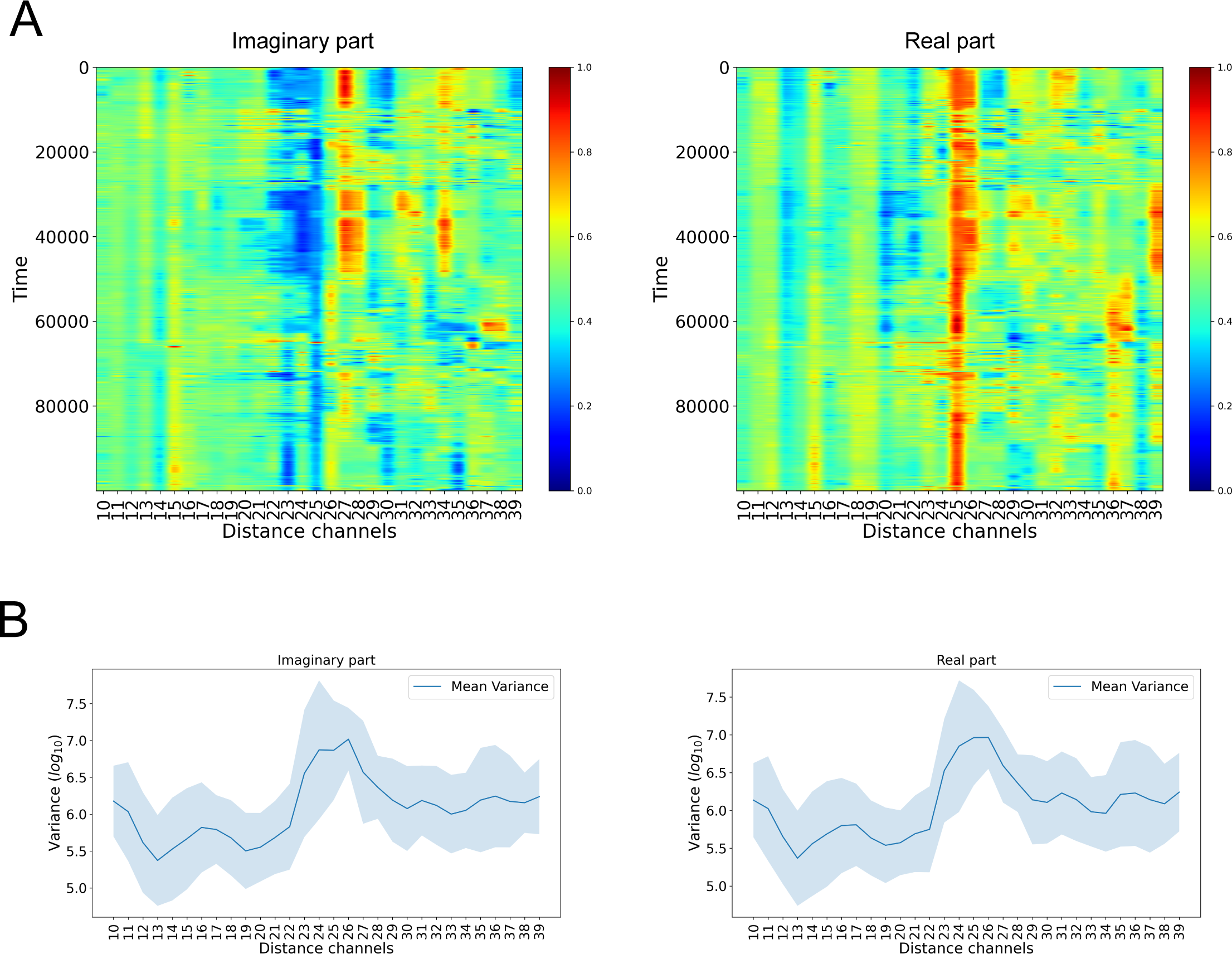
Heatmap visualization of mm-wave radar signals. (A) Example heatmaps of Imaginary and real components obtained from mm-wave radar. Each plot displays 30 channels in the distance direction, corresponding to 132 cm, and 100,000 data points (1,000 seconds) in the time direction. (B) The variance of mm-wave signals along the 30 distance channels.

**Figure S2.**
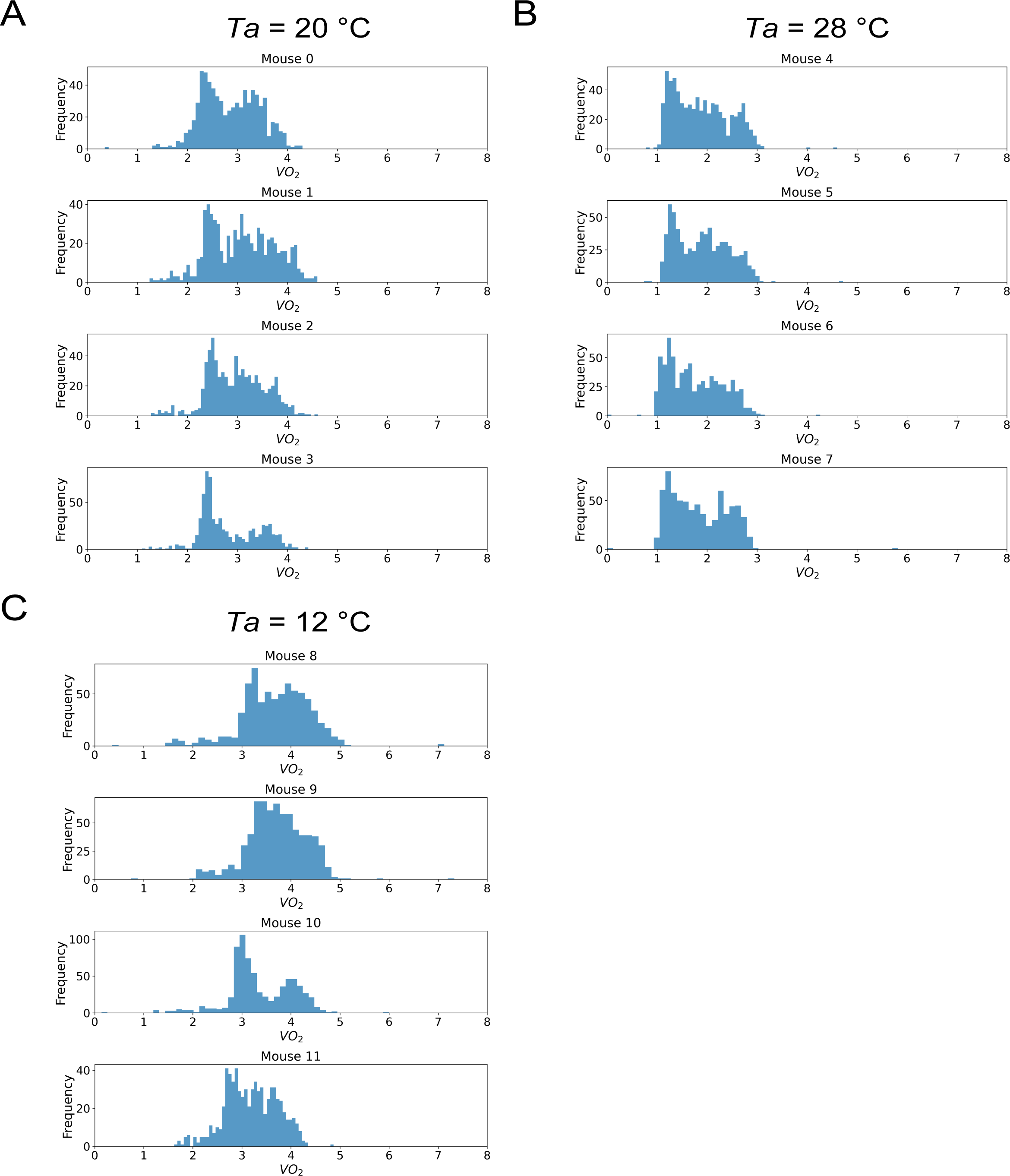
Distributions of oxygen consumption rates from each mouse. Distribution of oxygen consumption rates from each mouse under several *Ta*s (A: *Ta* = 20 °C, B: *Ta* = 28 °C, C: *Ta* = 12 °C).

**Figure S3.**
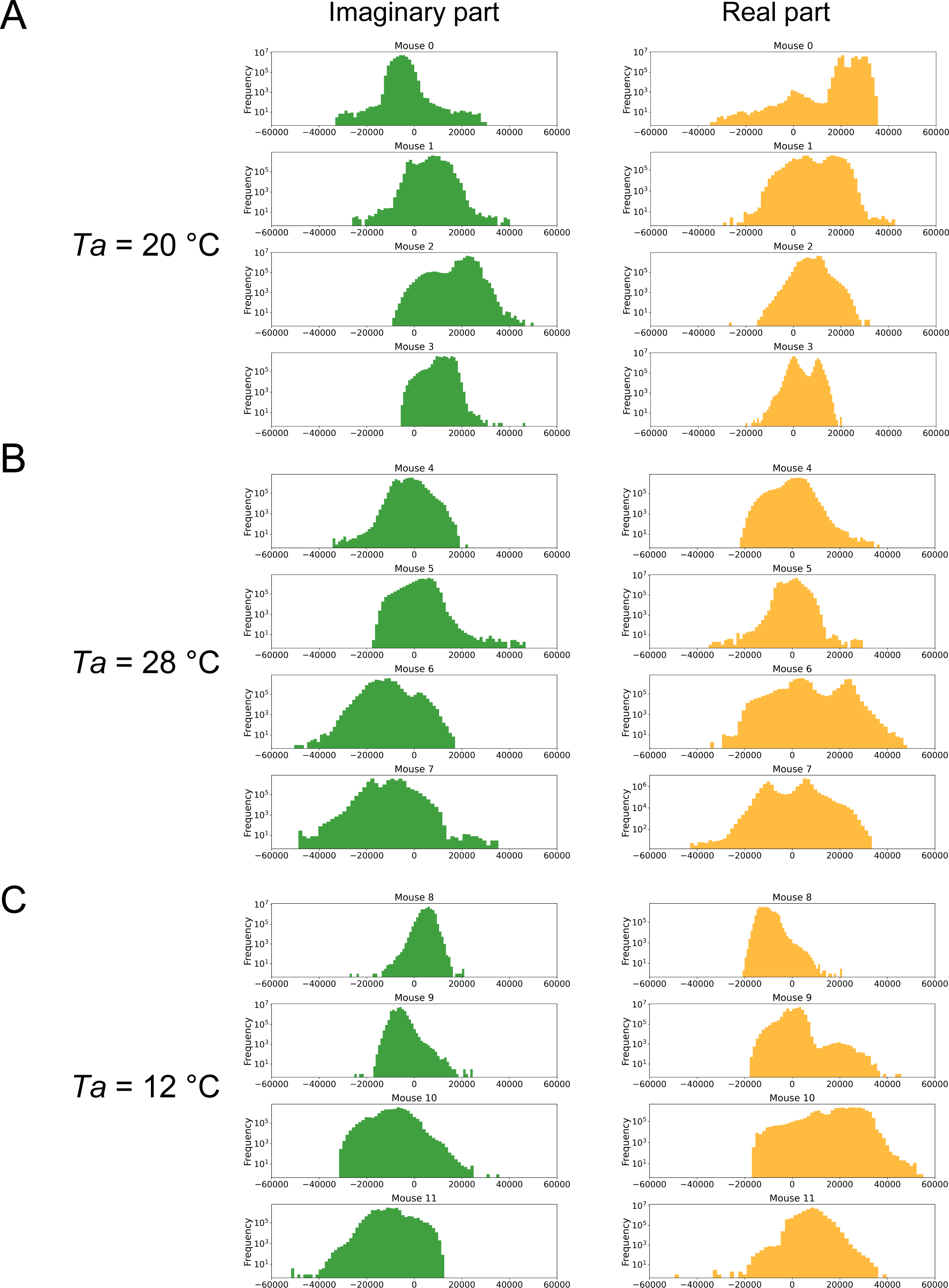
Distributions of mm-wave radar signals (I and Q) from each mouse. Distribution of mm-wave radar signals from each mouse under several *Ta*s (A: *Ta* = 20 °C, B: *Ta* = 28 °C, C: *Ta* = 12 °C). The green histogram depicts the imaginary component, while the orange histogram represents the real component.

**Figure S4.**
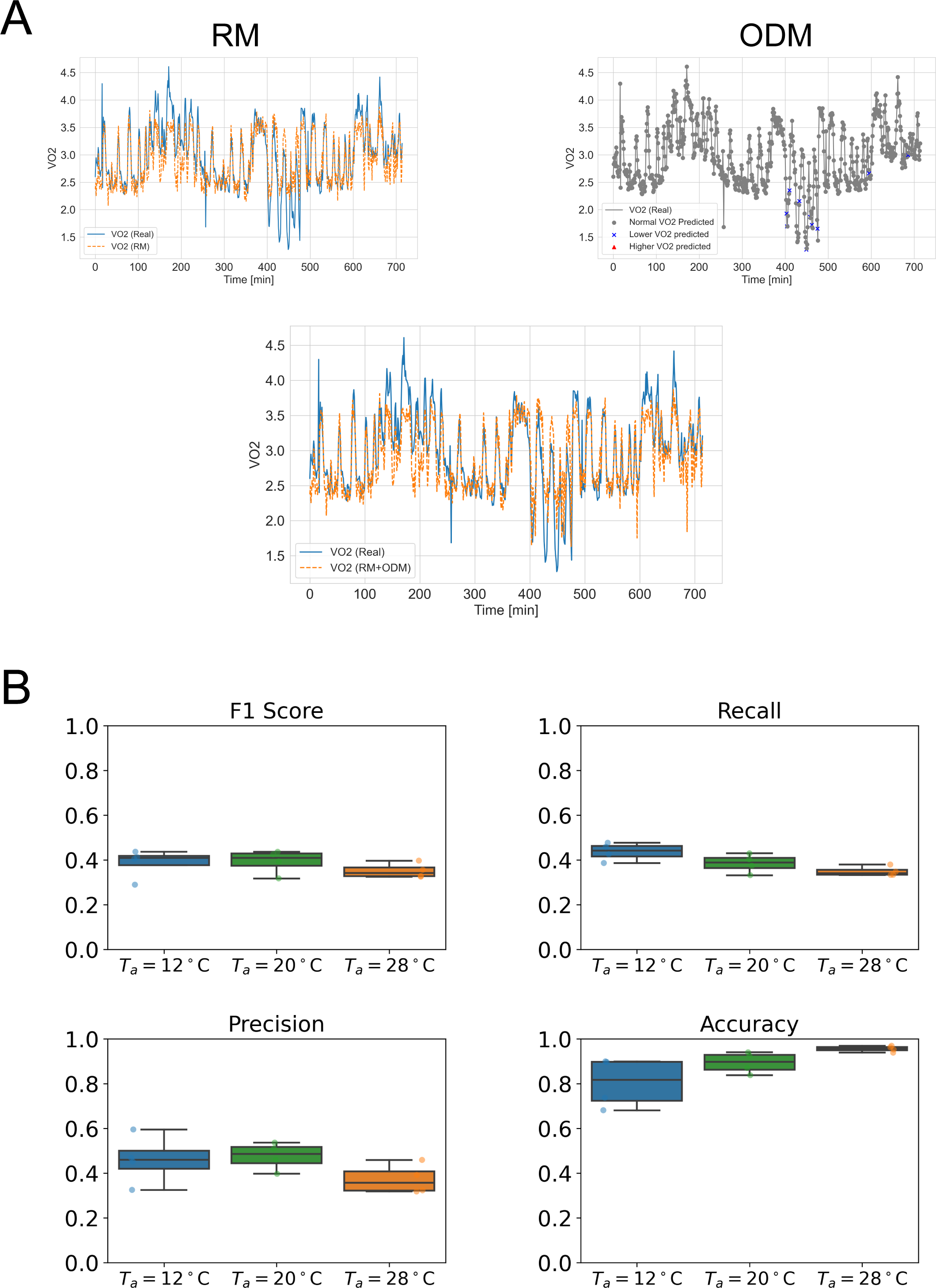
Performance of RMs. (A) Examples of outputs from the RM, ODM, and the final output (RM + ODM). The upper left panel displays the actual *VO_2_* (blue solid line) and the *VO_2_* predicted by the RM (orange dashed line). The upper right panel compares the actual *VO_2_* (gray solid line) with the categorizations made by the ODM, shown as ‘Normal’ *VO_2_* (gray circles), ‘Lower’ *VO_2_* (blue crosses), and ‘Higher’ *VO_2_* (red triangles). The bottom panel illustrates the actual *VO_2_* (blue dotted line) and the final output (RM + ODM; orange dashed line) obtained by linearly combining the RM and ODM outputs. (B) The performance metrics (F1 score, precision, accuracy, recall) of the ODM across different *Ta*s.

**Figure S5.**
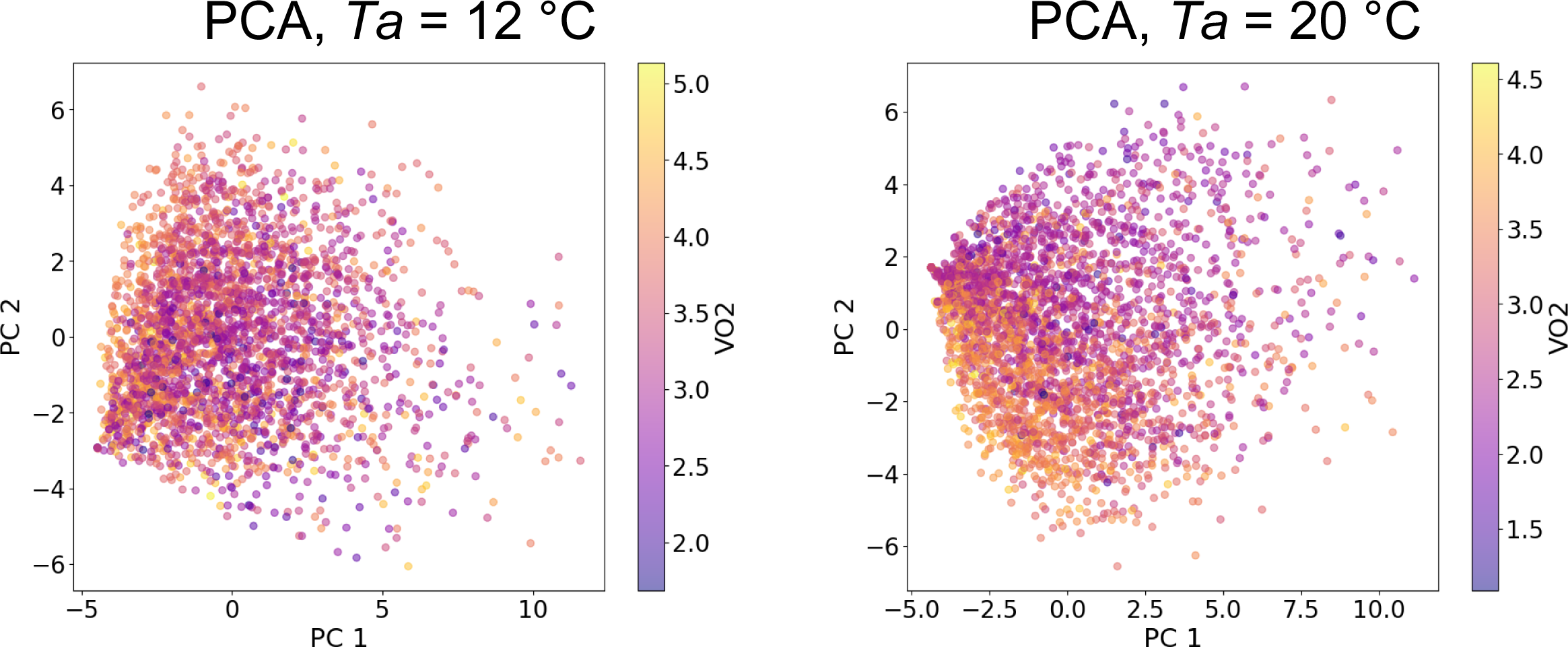
Results of PCA using mm-wave radar signals at *Ta* = 12 °C or *Ta* = 20 °C. PCA results at *Ta* = 12 °C (left panel) or *Ta* = 20 °C (right panel). Color gradients represent the *VO_2_* value at each point.

